# Efficient Phylogenetic Inference Using SNP-Based Approaches: A Comparison with Full Sequence Data

**DOI:** 10.1101/2025.03.10.642383

**Authors:** Vivekanantha Sanjeevan, Patrick König

## Abstract

Mutations in specific genomic regions or genes serve as reliable indicators of phylogenetic relationships, with single nucleotide polymorphisms (SNPs) playing a crucial role in population phylogenetic studies. Traditional distance-based phylogenetic algorithms have a time complexity proportional to l · n^2^, where n is the number of sequences and l is their length [1], [2]. This high computational cost becomes a bottleneck in phylogeny reconstruction, particularly when l > n.

To overcome this limitation, we propose an SNP-based approach to phylogenetic tree inference, focusing exclusively on variant (mutated) positions rather than entire sequences. This method significantly reduces computational time while maintaining accuracy. We compare phylogenies inferred from SNP data and full sequence data (including both SNPs and invariant sites) across multiple metrics.

Our results show that heuristic phylogenetic trees constructed from SNPs achieve parsimony scores nearly identical to those derived from full sequence data, as parsimony primarily depends on variant positions. Additionally, under the Jukes-Cantor 1969 (JC69) model, log-likelihood scores for SNP-based and full-sequence-based trees exhibit a strong correlation when evaluated using the same tree topology, branch lengths, and maximum likelihood parameters. These findings demonstrate that SNP-based methods can streamline phylogenetic analysis while preserving accuracy.

## I. Introduction

Advances in cost-effective sequencing technologies and increased computational power have revolutionized genomics, enabling researchers to generate and analyze vast amounts of high-quality genomic data [4]. This progress is evident in the exponential growth of sequenced species, exemplified by the Sanger Institute’s achievement of sequencing 243,633 human genomes in just 3.5 years [13]. Large-scale genomic projects like these highlight the growing need for efficient tools to analyze evolutionary relationships across genomes.

Phylogenetic trees, derived from DNA sequence data, provide critical insights into evolutionary relationships at various genomic levels [7]. However, the size of modern genomes—such as the 3.05 billion base pairs in the human genome—poses significant computational challenges, particularly when handling large datasets. Traditional methods for phylogenetic reconstruction often rely on calculating distance matrices, a process with a time complexity of *l · n*^2^ for *n* sequences of length *l* [1]. This becomes a bottleneck when both the sequence length (*l*) and the number of taxa (*n*) are large, as the computational complexity grows with both parameters.

Although genetic variation represents only about 0.1% of the genome, it introduces millions of differences across individuals. To manage this complexity efficiently, researchers often use variant call format (VCF) files, which store key information about genotypes and sequence variants [3]. By focusing on single nucleotide polymorphisms (SNPs), which represent mutated homologous positions, it is possible to construct phylogenetic trees while significantly reducing computational requirements without sacrificing essential information.

The choice of SNPs for phylogenetic analysis can introduce biases, particularly in regions with strong linkage disequilibrium (LD), where alleles at multiple loci tend to be inherited together [9]. This may result in an overrepresentation of certain genetic patterns, which can distort tree topology by exaggerating similarities between closely related taxa or generating misleading signals of divergence [10], [11]. To minimize these effects, it is crucial to carefully select SNPs and filter out loci with significant LD to ensure the phylogenetic tree remains both accurate and robust [9], [11].

To address these challenges, we developed a user-friendly bioinformatics pipeline that streamlines phylogenetic analysis by focusing exclusively on SNP data. This approach excludes non-contributory sequences, enabling the rapid construction of phylogenetic trees from VCF and FASTA files. The pipeline evaluates trees using parsimony and maximum likelihood scores, demonstrating a strong correlation between SNP-based models and those derived from full sequence data. This innovation provides an efficient and accurate solution for largescale phylogenetic studies.

## II. Methodology

We generate nucleotide sequence data in the International Union of Pure and Applied Chemistry (IUPAC) format for each sample from the Zarr dataset and compile it into a single FASTA file. Our analysis focuses on diploid genome sequences, processing the first and second alleles separately, and emphasizing single nucleotide polymorphisms (SNPs).

### A. Generating Nucleotide Sequences

For each sample, nucleotide sequences are generated by extracting reference allele characters, alternative allele characters, and genotype information from the Zarr dataset. The assignments are as follows:

- **Genotype 0/0:** Assign the reference allele to both allele sequences.
- **Genotype 1/1:** Assign the alternative allele to both allele sequences.
- **Genotype 0/1:** Assign the reference allele to the first allele sequence and the alternative allele to the second allele sequence.

The sequences for both alleles are stored in a NumPy array indexed by position.

### B. SNP Selection and Filtering

No filtering criteria (e.g., MAF thresholds, LD pruning, or quality-based selection) were applied. All SNPs were included to preserve full genetic variation, ensure reproducibility, and enable direct comparison between raw sequence-based and SNP-based phylogenetic trees. Future studies may incorporate filtering to refine phylogenetic resolution.

### C. Combining Alleles

To generate a single IUPAC nucleotide sequence, the first and second alleles are combined using IUPAC nucleotide codes. For example:

- **Sample 1:** Encodes `C` for homozygous reference alleles (0/0).
- **Sample 2:** Encodes `Y` (IUPAC for `C/T`) for heterozygous genotypes (0/1).
- **Sample 3:** Encodes T for homozygous alternative alleles (1/1).

For position 551^*st*^:

- Sample 1 encodes `C` (homozygous reference, 0/0).
- Sample 2 encodes `Y` (heterozygous, 0/1, `C/T`).
- Sample 3 encodes `T` (homozygous alternative, 1/1).
- sample 1: CGRCMG
- sample 2: YKGCCG
- sample 3: TGATCC

You can refer to the Python script available on GitHub: distance matrix.py on GitHub

### D. Sequence Data Generation Using Reference FASTA

We use SAMtools and BCFtools [14], [15] to process sequencing data, including sorting, querying, and variant calling. To generate sequence data, we combine VCF files with reference FASTA files. The required inputs are:

- VCF file and reference FASTA file.
- Chromosome ID, start position, and end position for the target region.
- Names of selected samples and alleles from the FORMAT/GT field.

The sequences are stored in a FASTA file, where each entry is formatted as follows:

- Line 1: Identifier starting with >.
- Line 2: Sequence data in IUPAC or nucleotide codes.

### E. Computation of Pairwise Distance and Hierarchical Clustering

We compute the pairwise Hamming distance between sequences using a custom function. The distance matrix is then generated, organizing pairwise distances into a square matrix where rows and columns represent sample names.

Heuristic hierarchical clustering is applied to merge clusters iteratively based on average linkage criteria:

### F. Average Linkage Algorithm

The clustering procedure is summarized in the following algorithm:

#### Algorithm 1

Linkage Algorithm

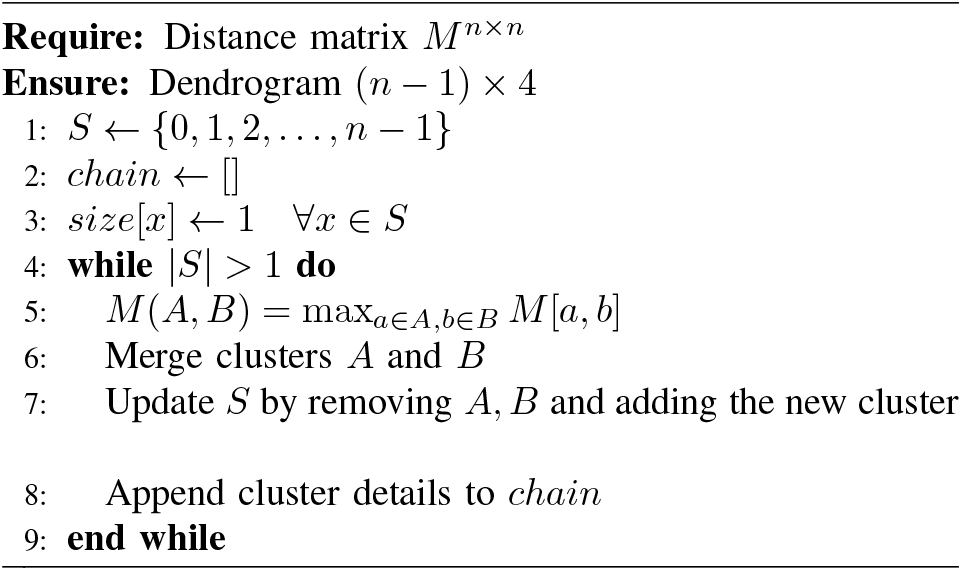

## PHYLOGENETIC TREE ANALYSIS AND RESULTS

We analyze the Barley Morex V2 pseudomolecule dataset, which includes 300 samples and 223,387,147 SNPs, to evaluate phylogenetic tree construction using the average linkage clustering algorithm. For this analysis, we focus on the following regions of Chromosome 1, considering varying sequence lengths:

- Chromosome 1: 1–1,000
- Chromosome 1: 1–10,000
- Chromosome 1: 1–100,000
- Chromosome 1: 1–250,000
- Chromosome 1: 1–500,000
- Chromosome 1: 1–750,000
- Chromosome 1: 1–1,000,000

## III. Comparison OF ParsimonyScores BETWEEN SNPS AND Full Sequence Data

Figure 1 illustrates the phylogenetic trees derived from SNP and full sequence data for four species using the Fitch algorithm. Subfigure 1(a) shows the tree constructed using SNP data, with a parsimony score of 2. Subfigure 1(b) represents the tree derived from full sequence data, also with a parsimony score of 2. For homologous positions 2 to 6, the characters are identical across all species, resulting in a parsimony score of 0 when calculated for each homologous position individually, as no character changes occur at these positions.

**Fig. 1:**
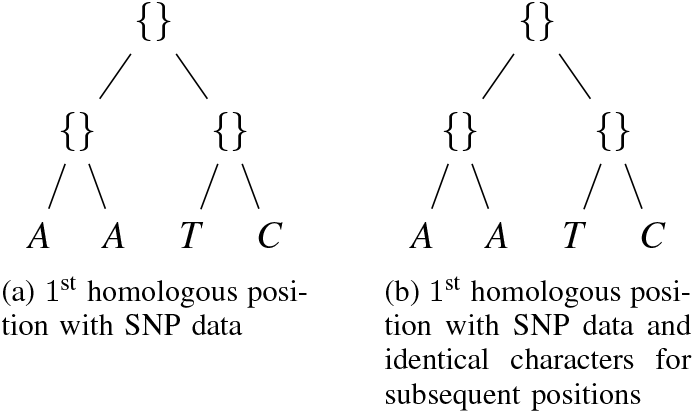
Example of SNP (a) and Sequence (b) data using the Fitch algorithm

Let *T* represent a specific tree topology, potentially selected randomly from the set of all possible trees. The parsimony score *L*(*T*) of tree *T* is defined as:

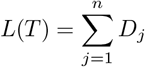

where *D*_*j*_ is the parsimony score for the *j*^th^ homologous position, and *n* is the total number of sites (or characters) in the alignment. Parsimony is computed for each homologous position, and the total parsimony score is the sum of the parsimony scores across all homologous positions, as defined by Felsenstein (2004) [27, p. 270].

Table II compares parsimony scores between SNP and full sequence data for phylogenetic trees constructed using a genotype-based distance matrix and the average linkage algorithm. The results indicate that SNP data yields nearly identical parsimony scores compared to full sequence data, demonstrating its efficiency in phylogenetic tree computation.

**TABLE I:**
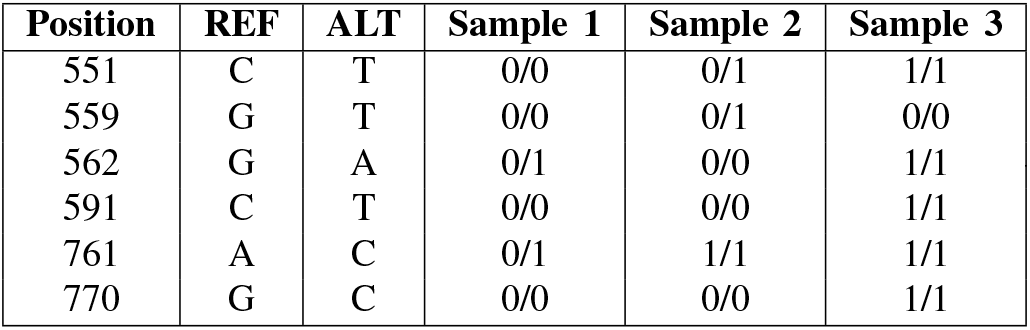
Example of VCF file.

**TABLE II:** Comparison of Parsimony Scores between SNP and Full Sequence Data.

| Sequence Length | Parsimony Score |  |
| --- | --- | --- |
|  | SNP Data | Full Sequence Data |
| 1,000 | 30 | 30 |
| 10,000 | 53 | 42 |
| 100,000 | 11,822 | 11,821 |
| 250,000 | 87,335 | 87,335 |
| 500,000 | 166,431 | 166,419 |
| 750,000 | 318,403 | 318,392 |
| 1,000,000 | 365,517 | 365,517 |

We construct the genotype data-based distance matrix of the phylogenetic tree using the average linkage algorithm and then evaluate the parsimony score of the phylogenetic tree using SNP and full sequence data. Table II shows that both data models computed almost the same score on the genotype distance matrix of the phylogenetic tree. Non-mutated homologous positions of all samples do not affect the parsimony score. Therefore, we can use the SNP data rather than sequence data to compute the parsimony of the phylogenetic tree.

## IV. Comparison OF Likelihood Scores BETWEEN SNP AND Full Sequence Data using the JC Model

We use the simple JC69 model to compare the loglikelihood of SNP and full sequence data. The JC69 model was chosen for its simplicity and because it assumes equal substitution rates between all nucleotide pairs [20]. While more complex models like GTR allow for unequal rates, they require more parameters and can overfit, especially with small datasets [21], [22]. The JC69 model is effective for analyzing log-likelihood scores without added complexity, making it ideal for large datasets where computational efficiency and interpretability are crucial [23].

The **Jukes-Cantor (JC) model** assumes equal substitution rates between all nucleotides. It is based on a continuous-time Markov process, where the rate matrix *Q* governs nucleotide transitions. The probability of transition over time *t* is given by:

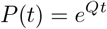

The JC rate matrix is:

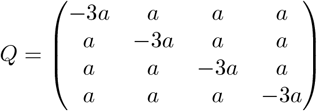

For a given phylogenetic tree, the likelihood is computed as:

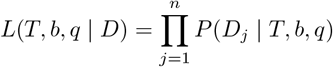

To avoid numerical underflow, the logarithmic likelihood is used:

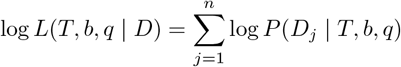

Figure 2(b) shows that the log-likelihood score of the SNP data is generally higher compared to full sequence data. This is because full sequence data contains both SNP and non-SNP data, while the VCF file contains only SNP data. SNP data typically leads to a higher likelihood due to the absence of invariant positions, which are not informative. However, the relationship between the log-likelihood scores of SNP and full sequence data depends on the composition and mutation patterns in the data.

**Fig. 2:**
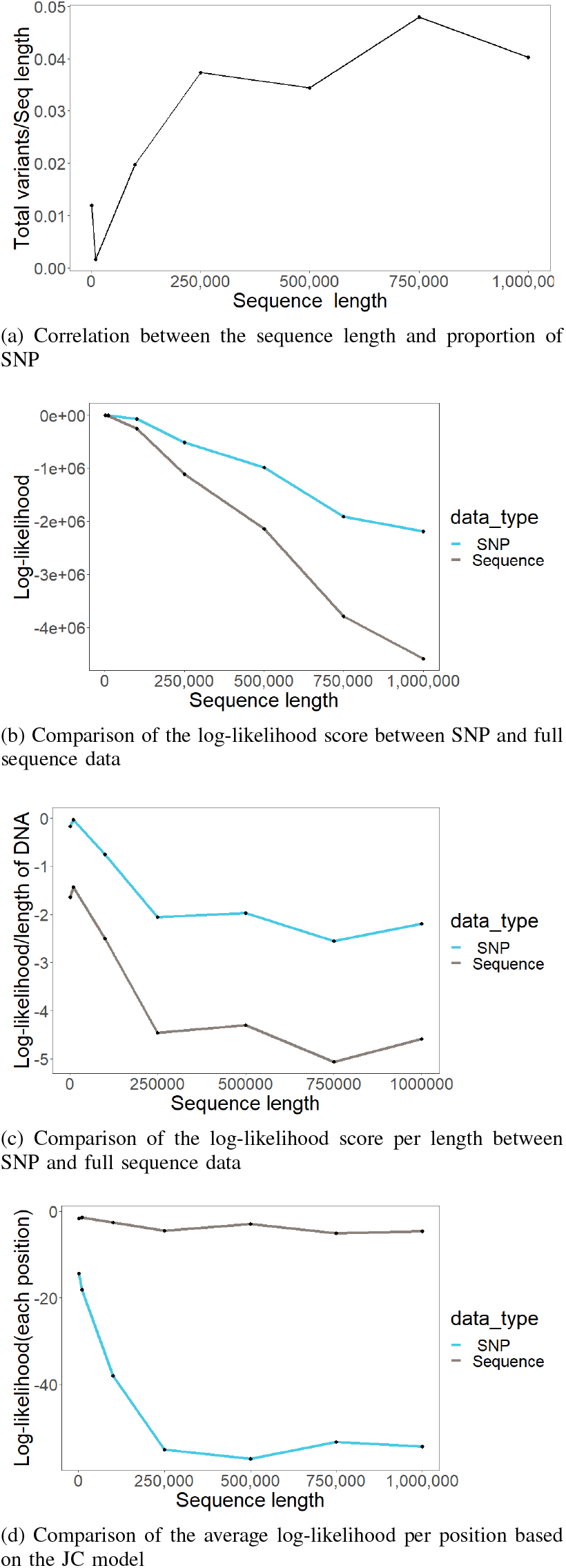
Comparison of the log-likelihood between SNP and full sequence data (b-d) with respect to the proportion of SNP (a)

**Fig. 3:**
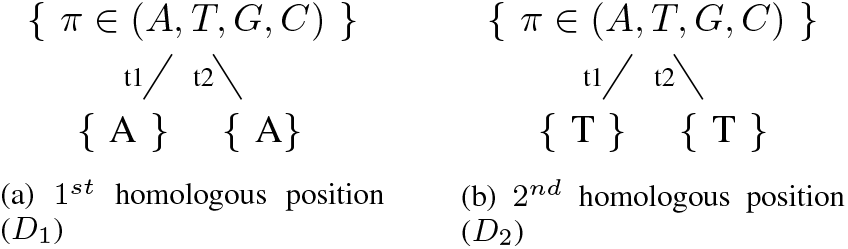
Example of the *i*^*th*^ homologous position of data *D*

The log-likelihood of each homologous position can produce negative log-likelihoods, and as the number of homologous positions increases, the overall negative log-likelihood generally becomes more pronounced. This occurs because additional positions provide more opportunities for mutations, which influence the likelihood calculation.

There is a correlation between the log-likelihood of SNP and full sequence data (Figure 2(c)), which depends on the proportion of SNP or variant data (Figure 2(a)). As the proportion of variants increases, the log-likelihood of SNP data becomes more distinct from full sequence data, due to the exclusive consideration of mutated nucleotides in SNP data.

The average log-likelihood score per character of SNP data tends to be lower compared to full sequence data (Figure 2(d)) because SNP data only considers mutated nucleotides. Since the probability of a nucleotide changing into another is typically very low, the log-likelihood per character for SNP data might be lower when compared to the full sequence data, which includes both invariant and variant sites.

Consider the following two DNA sequences that contain the same characters in each homologous position:

- Sample 1: AT
- Sample 2: AT

Matrix 1 explains the rate matrix of the JC model, where the mutation rate between two different nucleotides is always the same. Each diagonal element is computed as the sum of the rest of the row multiplied by -1.

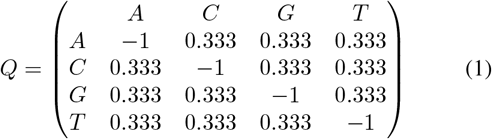

Any given nucleotide change to another nucleotide does not happen in discrete time. Instead, it occurs in continuous time, and we must compute the probability that, for an infinitesimally small period *t*, the change occurs. The continuous-time Markov chain transition-probability matrix is computed as *P* (*t*) = *e*^*Q*\**t*^, where *Q* is the predefined rate matrix (1) for the JC model.

Matrix 2 explains the continuous-time Markov chain of the rate matrix with respect to assumption time 1 *×* 10^−6^ for the JC model, computed as *P* (*t*) = *e*^*Q*\*1*×*10−6^. The probability that each nucleotide remains the same without change is higher than the probability of it changing to another nucleotide during evolution.

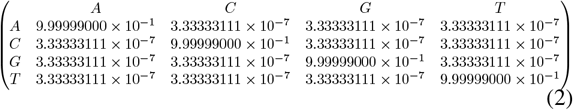

For the given trees in Figure 3(a) and the continuous time transition rate matrix 2, we compute the log-likelihood score of the 1^*st*^ homologous position data (*D*_1_) as follows:

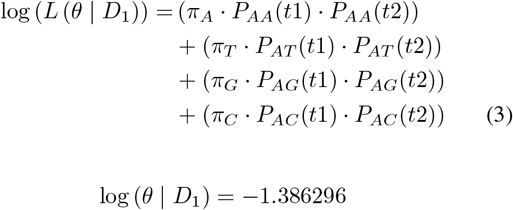

We can apply the same computation to the 2^*nd*^ homologous position of the given samples, and the results will produce the same log-likelihood score as the 1^*st*^ homologous position if the characters of the two samples are identical. The log-likelihood of the entire sequence can be calculated by adding the log-likelihood values of each position.

For example, if we compute the log-likelihood for the 2^*nd*^ homologous position of the above samples, the value would be

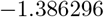

and the sum of the log-likelihoods for the sequence would be

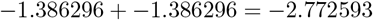

Consider the following DNA sequences that contain different characters in each homologous position:

- Sample 1: AC
- Sample 2: GT

For the trees in Figure 4(a) and the continuous time transition rate matrix 2, we compute the log-likelihood score of the 1^*st*^ homologous position data (*D*_1_) as:

**Fig. 4:**
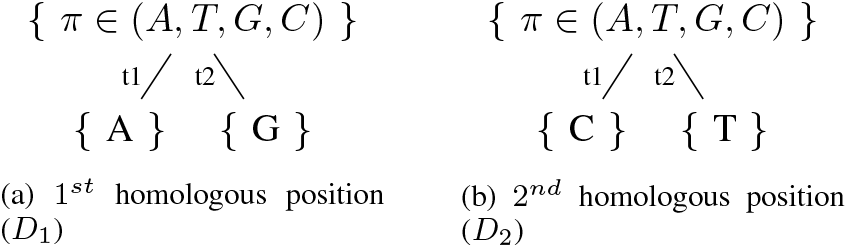
Example of the 1^*st*^ homologous position of two samples in the tree with four possible nucleotides appearing at the root, and *t*_1_, *t*_2_ are the branch lengths with respect to the edges

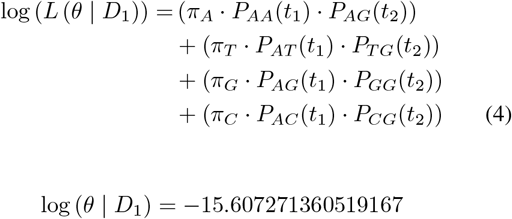

Similarly, we compute the log-likelihood score for the 2^*nd*^ homologous position of the samples, and if the characters are different, the log-likelihood score will be the same as for the 1^*st*^ homologous position. The log-likelihood score of the entire sequence is the sum of the log-likelihoods for each position.

For instance, if the log-likelihood for the 2^*nd*^ homologous position of the samples is

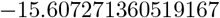

then the sum of the log-likelihoods for the sequence is

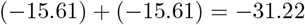

Figure 5 illustrates the two phylogenetic trees inferred from SNP and full sequence data. Specifically, Figure 5(a) presents the tree constructed from SNP data, while Figure 5(b) shows the tree derived from sequence data. Assuming a branch length of 1 *×* 10^−6^, the log-likelihood score based on SNP data is

**Fig. 5:**
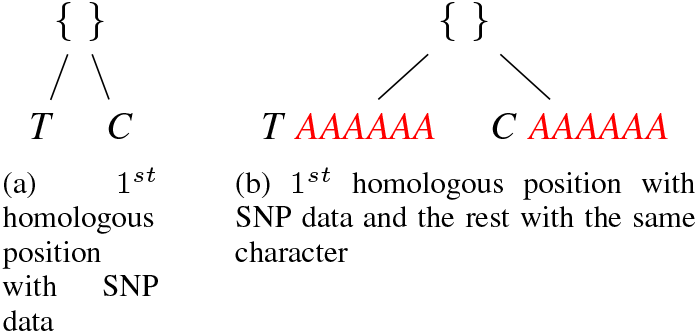
Example of SNP (a) and Sequence (b) for loglikelihood computation

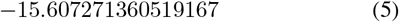

whereas the log-likelihood score for the sequence data is given by

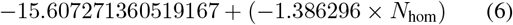

where *N*_hom_ represents the number of homologous positions with the same character.

Conclusion

In this study, we compared phylogenetic tree models constructed from full genomic sequences and SNP data, evaluating their performance using the Fitch parsimony algorithm and the Jukes-Cantor (JC) model. Our results indicate that both data types yield comparable parsimony scores, suggesting that SNP data is a reliable alternative for phylogenetic analysis. Additionally, SNP data generally produces higher loglikelihood scores than full sequence data, likely due to the exclusion of invariant positions, which simplifies analysis and improves model fit. As the proportion of SNPs increases, the log-likelihood scores for SNP and full sequence data diverge further, highlighting the advantage of SNP-based methods in highly variable genomic regions. Despite these differences, the log-likelihood scores of both data types remain correlated, suggesting that SNP and full sequence data provide consistent evolutionary insights.However, it is important to note that while SNP-based methods offer computational efficiency, they come with trade-offs in terms of accuracy. The exclusion of invariant positions may simplify the analysis but also limits the amount of information captured from the full genomic sequence.

The phylogenetic trees generated by our pipeline have broad applications, including tracking evolutionary trajectories of pathogens, understanding population structure, and identifying functionally significant genetic variants. For example, in oncology research, these trees could help reconstruct clonal evolution in tumors, offering insights into treatment resistance mechanisms. These potential applications highlight the versatility of our pipeline and its value beyond basic phylogenetic analysis.

Overall, our findings suggest that while SNP data offers a computationally efficient and reliable alternative to full genomic sequences, the trade-off between computational speed and accuracy must be carefully considered, particularly for studies requiring a high degree of precision. This is especially relevant in large-scale studies where processing time and computational resources are critical factors.

## V. Code AND Data Availability

The code for the phylogenetic analysis pipeline is available on GitHub in two repositories:

- [Phylogenetic Statistics Repository](https://github.com/Sanjeevan-informatik/Phylogenetic_Statistics)
- [VCF to Phylogenetic Repository](https://github.com/Sanjeevan-informatik/VCFToPhylogenetic)

The data used for this study is the Barley Pseudomolecules Morex v2.0 (2019). The dataset consists of 8 pseudomolecules, with the largest scaffold measuring 675,310,294 bp and an average sequence size of 542,842,368 bp. The L50 for this dataset is 624,247,919 bp. For further details, refer to the publication doi:10.1101/631648. The dataset is available for download at the eDal repository.

## APPENDIX

### Proofs of Theorems

#### Lemma 1.

*Let T be a phylogenetic tree, and let A represent a set of aligned sequences from a given set of samples. Suppose we partition A into two disjoint subsets:*

- *B, the set of sequence positions corresponding to single nucleotide polymorphisms (SNPs), and*
- *C, the set of sequence positions corresponding to invariant (non-SNP) sites*.

*If each homologous position evolves independently, then applying the Fitch algorithm to the full sequence data is equivalent to applying it only to the SNP subset, i*.*e*.,

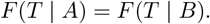

*Proof*. The Fitch algorithm computes the parsimony score *F* (*T* | *A*) as the sum of individual site contributions:

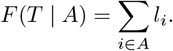

Since *A* = *B* ∪ *C*, we rewrite this sum as

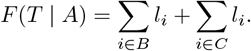

For a given site, the Fitch algorithm assigns a cost only when a character state change occurs. Because all sites in *C* are invariant, they do not contribute to the total cost, implying

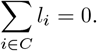

Thus,

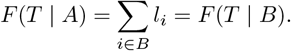

#### Lemma 2.

*Let T be a phylogenetic tree with branch lengths b and a transition rate matrix q (e*.*g*., *JC69 or K2P). Suppose the sequence data A is partitioned into SNP sites B and non-SNP sites C, such that A* = *B*∪ *C. If each homologous position evolves independently, then the log-likelihood computed from SNPs provides a lower bound on the log-likelihood of the full sequence data:*

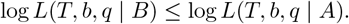

*Proof*. The total log-likelihood for sequence data *A* given tree *T* and parameters (*b, q*) is computed as the sum over all sites:

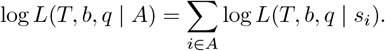

Since *A* = *B* ∪ *C*, we separate the terms:

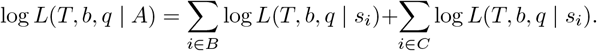

For non-SNP sites *C*, the likelihood contributions primarily come from stationary base frequencies *π*, leading to

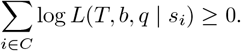

Thus,

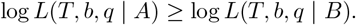

